# Neural dynamics of predictive timing and motor engagement in music listening

**DOI:** 10.1101/2023.04.29.538799

**Authors:** Arnaud Zalta, Edward W. Large, Daniele Schön, Benjamin Morillon

## Abstract

Why do humans spontaneously dance to music? To test the hypothesis that motor dynamics reflect predictive timing during music listening, we built melodies with varying degrees of rhythmic predictability. Magnetoencephalography data showed that while auditory regions track the rhythm of melodies, intrinsic neural dynamics at delta (1.4 Hz) and beta (20-30 Hz) rates in the dorsal auditory pathway embody the experience of groove. Critically, neural dynamics are organized along this pathway in a spectral gradient, with the left sensorimotor cortex acting as a hub coordinating groove-related delta and beta activity. Combined with predictions of a neurodynamic model, this indicate that spontaneous motor engagement during music listening is a manifestation of predictive timing effected by interaction of neural dynamics along the dorsal auditory pathway.

**One-Sentence Summary:** Interacting neural dynamics along the dorsal auditory pathway effect the experience of groove during music listening.

## Introduction

Dancing to the beat of music is universal (*1, 2*). Music is most often considered as an auditory phenomenon, but under an ecological and phylogenetic perspective it is tightly coupled to dance (*3, 4*). Dance requires synchronizing body movements with the musical rhythm via audio-motor interactions (*5–8*). Not all music induce dance equally, but why some urge us to dance? What are the brain mechanisms supporting the musical wanting-to-move experience called groove (*9–15*)?

Combining a neurodynamic model (*16, 17*) with the concept of auditory active sensing (*18, 19*) provides a canonical dynamical framework to understand the spontaneous emergence of movements during music listening. Neurodynamic models offer a computationally rigorous and neurophysiologically plausible approach to understanding the emergence of music-related cognitive phenomena (*16*). Active sensing (*20–23*) refers to the fact that perception is strongly shaped by motor activity, which notably imposes temporal constraints on the sampling of sensory information, in particular in the delta band, the range of natural movements and of music (*24*). During auditory perception, the motor system is recruited through passive listening of temporally structured musical rhythms (*25, 26*), encodes temporal predictions information and can optimize auditory processing (*7, 19, 27–29*). Here, we hypothesize that manipulating the rhythmic properties of music suffice to induce a covert motor engagement during music listening, via changes in audio-motor neural dynamics.

## Results

We created a stimulus set of 12 short melodies with a 2 Hz beat. To vary their level of rhythmic predictability, three variants were derived from each melody, using an ascending degree of syncopation (low, medium, high; Fig. 1a-c). As expected, the degree of syncopation was inversely proportional to the amplitude of the acoustic dynamics at 2 Hz (r^2^(34) = 0.81; *p* < 0.001; Fig. 1d). Nonetheless, when asked to reproduce the rhythm of their dance step while listening to the melodies, participants (n = 15) predominantly moved at the 2 Hz beat across conditions (Fig. 1e). Next, we recorded magnetoencephalography (MEG) data while participants (n = 29) listened to the melodies. A first analysis showed an absence of 1:1 mapping between the acoustic temporal envelope and neural frequencies. Instead, acoustic dynamics are encoded at 2 Hz and in a lesser extent at its harmonics (q < 0.05, FDR-corrected; Fig. 1f; see Methods). Thus, both behavioral motor and neural cortical dynamics principally track the 2 Hz beat.

**Figure 1.**
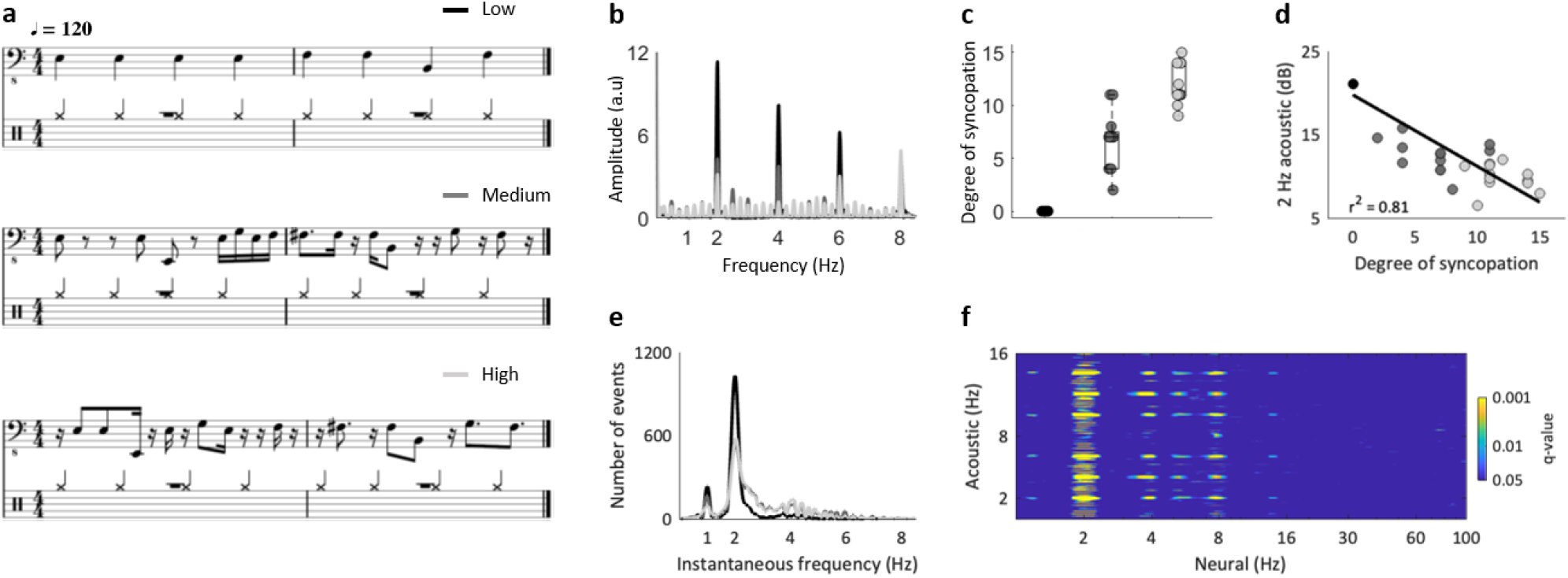
Stimulus set. Twelve 8-seconds melodies with a 2 Hz beat were created. For each melody, three variants were designed to vary the level of rhythmic predictability (degree of syncopation) while minimizing other acoustic variations. **(a)** Example of a melody with a low (black), medium (grey) or high (light grey) degree of syncopation. **(b)** Averaged modulation spectrum of the acoustic temporal envelope of the melodies, for each of the three conditions. **(c)** Degree of syncopation of the melodies, grouped by condition. Each dot represents one melody. **(d)** Amplitude of the acoustic envelope at 2Hz (in dB; ‘*2 Hz acoustic*’), as a function of the degree of syncopation, across melodies. Data were approximated with a linear function. Pearson’s r-squared is reported. Shades of grey indicate the conditions. **(e)** Behavioral tapping experiment: distribution of the instantaneous frequency of finger tapping per condition, cumulated across melodies and participants, recorded while participants were reproducing the rhythm of their dance step while listening to the melodies. **(f)** MEG experiment: Statistical map of neural coding of the acoustic temporal modulation spectrum, from the power spectrum of the whole-brain MEG signals recorded while participants were listening to melodies (q < 0.05, FDR-corrected).

We also asked participants, either online (n = 66) or during MEG recordings, to rate groove for each melody (Fig. 2a). Participants’ desire to move highly correlates with the degree of syncopation, but in a nonlinear manner (Fig. 2b-c; see Supplementary Results). This inverse U-shape profile is well approximated with a quadratic function (online: adjusted r^2^(33) = 0.73; MEG exp.: adjusted r^2^(33) = 0.67), confirming previous findings of moderately syncopated melodies inducing strongest wanting-to-move experiences (*12, 14, 15, 30*). This behavior shows that motor engagement neither indexes temporal predictability (highest in the low-syncopated condition), nor temporal prediction errors (highest in the high-syncopated condition) but is compatible with the notion of a precision-weighted temporal prediction error computation (*5, 31*).

**Figure 2.**
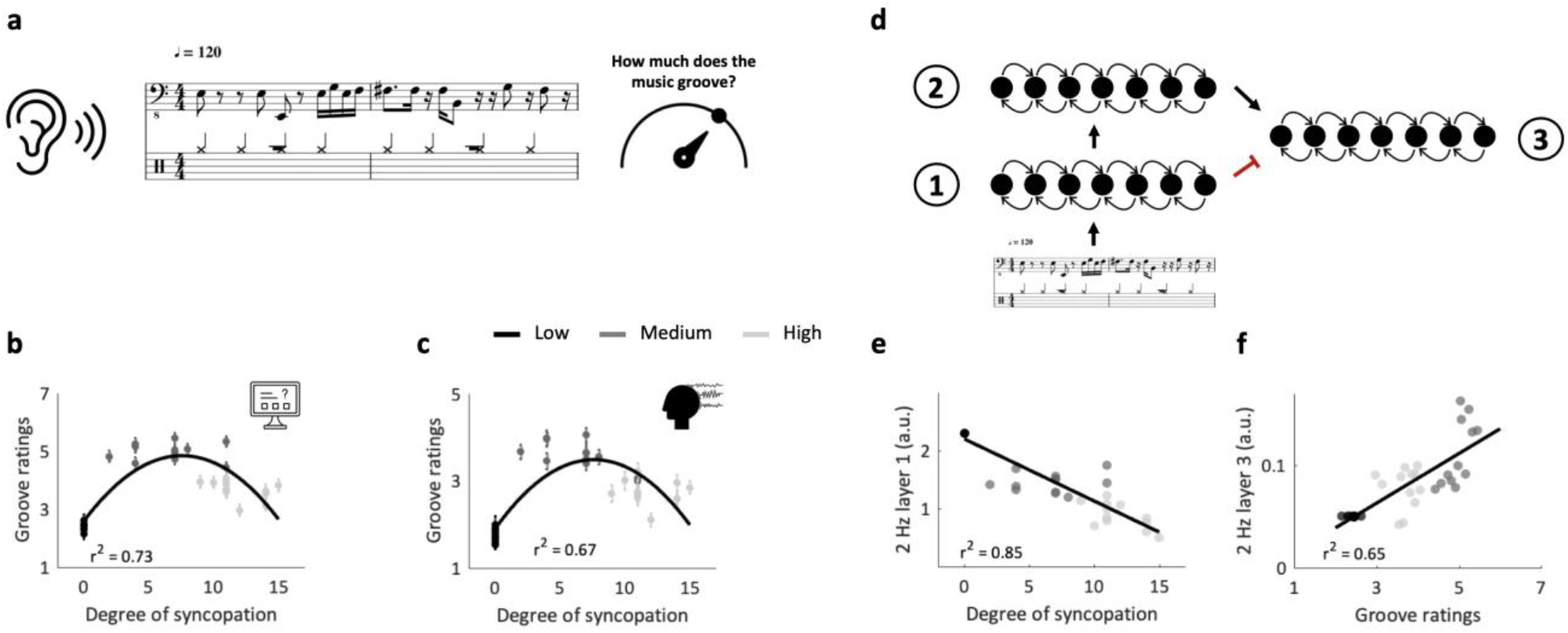
Behavioral experiments and neurodynamic model. **(a)** Main experimental design: in two experiments (online and MEG), participants listened to melodies. After each melody, they were asked to rate it in terms of groove, defined as the extent to which they wanted to move to the music. **(b-c)** Behavioral rating of participants (groove) acquired **(b)** online (from 1 to 7) or **(c)** during the MEG experiment (from 1 to 5), as a function of the degree of syncopation, across melodies. Data were approximated with a quadratic function. Adjusted r-squared is reported. Shades of grey indicate the conditions. Error bars indicate SEM. **(d)** Neurodynamic model: Each layer represents a network of coupled oscillators at different frequencies. The rhythm of melodies was input in the first layer. Arrows represent the coupling across layers (black is excitatory, red is inhibitory). **(e)** Amplitude of the output of layer 1 at 2 Hz, as a function of the degree of syncopation, across melodies. Data were approximated with a linear function. Pearson’s r-squared is reported. Shades of grey indicate the conditions. **(f)** Amplitude of the output of layer 3 at 2 Hz, as a function of groove ratings (from the online experiment), across melodies. Same conventions as in (e).

Next, we created a neural network model composed of three layers, each layer representing a network of oscillators spanning a range of frequencies (*16, 32, 33*). Following previous modelling work on the perception of rhythmic pulse (*16*), the melodies’ rhythms were presented as input to a first network layer (layer 1), which modelled auditory cortical dynamics as an oscillatory network operating near a Hopf bifurcation, and two other network layers (layers 2 and 3) which modelled motor cortical dynamics operating near a double limit cycle bifurcation (Fig. 2d; see Methods). This 3-layer neurodynamic model accounts for the nonlinear transformation from a syncopated stimulus rhythm to the subjective experience of groove. Indeed, we observed a dissociation between 1-a strong linear correlation of the degree of syncopation with the 2Hz activity in layer 1 (r^2^(34) = 0.85; *p* < 0.001; Fig. 2e) and 2-a strong linear correlation of groove ratings with the 2Hz activity in layer 3 (online experiment: r^2^(34) = 0.66; *p* < 0.001; Fig. 2f), with far less contributions of the other layers (see Supplementary Results and Fig. S1a).

We then interpreted our model to understand the mechanisms that may underlie the experience of groove. The first two network layers were intended to reflect the auditory rhythm (auditory cortex) and the perception of pulse and meter (motor planning cortex) (*16*). Oscillations in layer 2 may be considered as temporal predictions, i.e., *expectations* about the timing of rhythmic events (*34, 35*). Critically, layer 2 resonances arise not based on a learned model but based on mode-locking of neural oscillators to other frequencies in the rhythm. To these, we added a third layer that received inhibitory connections from layer 2 and excitatory connections from layer 1 (see Methods). Hence, layer 3 responds to the *difference* between the time-dependent oscillations of layer 2 (pulse/meter) and layer 1 (auditory rhythm) and can thus be interpreted as reflecting the divergence between temporal predictions (layer 2) and the actual input (layer 1). That groove ratings only correlate with amplitude in layer 3 indicates that the groove phenomenon may be described in terms of the timing of auditory events (layer 1) relative to *expectancies* (layer 2) embodied in oscillations.

We next analyzed the neural dynamics of cortical activity while listening to the melodies. We first estimated the 1/f-rectified power spectrum (1-100 Hz; see Methods) of neural activity at the source-level and observed that the spatial and spectral dimensions are closely related, in the form of a bilateral spectral gradient along the dorsal auditory pathways (Fig. 3a). The frequency of the dominant activity progressively increases from auditory regions (< 10Hz) to the motor cortex (20-30 Hz), up to the inferior frontal cortex (> 30 Hz; activity > 45 Hz was not observed).

**Figure 3.**
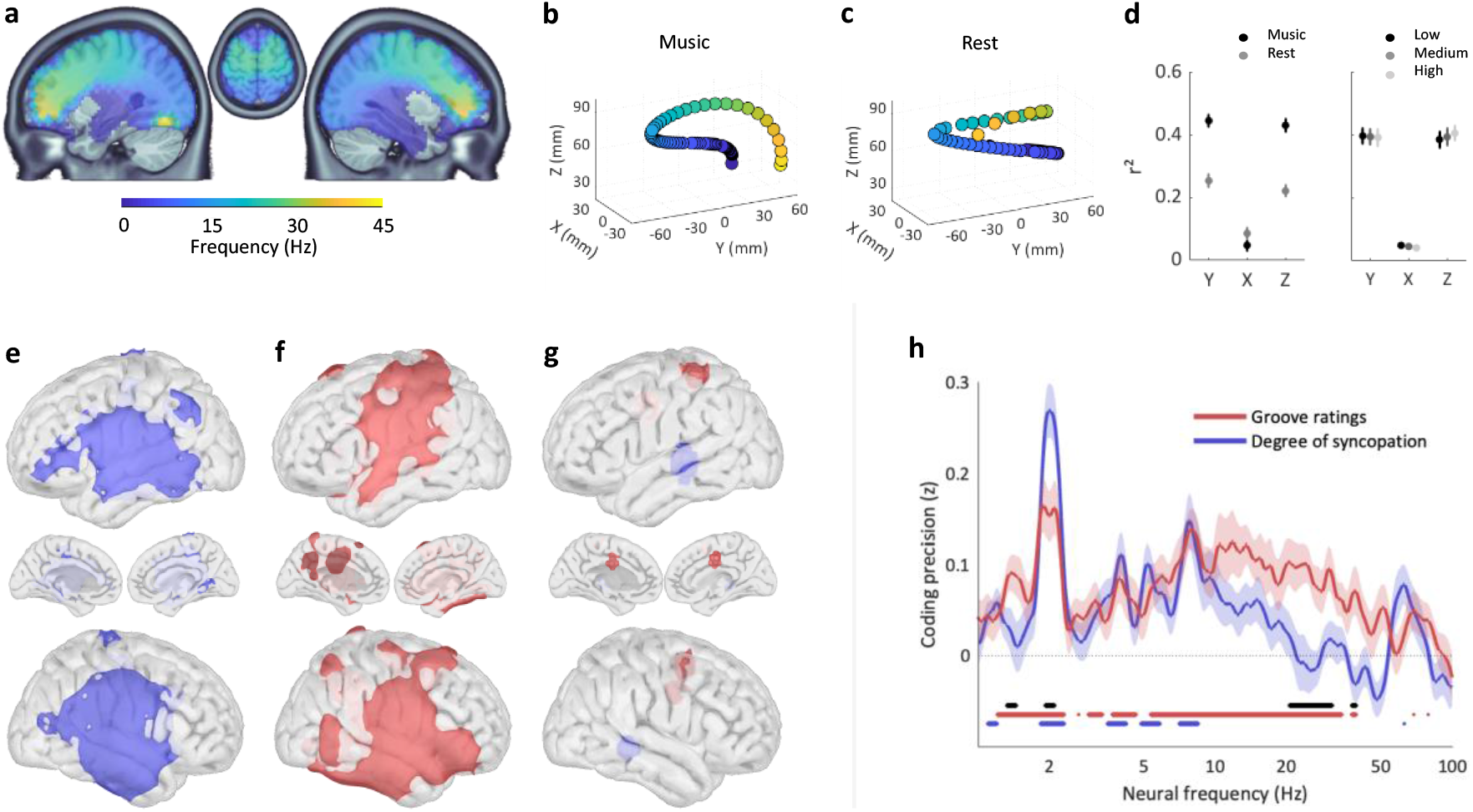
MEG experiment: Spatial and spectral coding of degree of syncopation and groove ratings. **(a)** Dominant frequency across the brain volume during music listening of the melodies (1-45 Hz; after removal of the 1/f decay of the neural power spectrum). **(b)** Data were approximated at the group-level with a polynomial function, independently for each dimension (X, Y, Z) of the MNI space. **(c)** Control analysis: Same as in (b) but from a resting-state MEG dataset. **(d)** Comparison of the quality of fits (r^2^), estimated at the individual level, between (left) the music and rest MEG datasets and (right) the three melodic conditions. Error bars indicate SEM. **(e-f)** Spatial map of neural coding of **(e)** the degree of syncopation (blue) and **(f)** groove ratings (red), from the power spectrum (1-100 Hz) of neural data recorded while participants were listening to melodies. Results reported at q < 0.005, FDR-corrected. **(g)** Spatial map of the difference in coding precision between degree of syncopation and groove ratings (p < 0.005, uncorrected). **(h)** Spectrum of neural coding of degree of syncopation (blue) and groove ratings (blue), from whole-brain MEG signals. Red and blue lines indicate frequencies with significant coding values (q < 0.005, FDR-corrected). The black line indicates frequencies with significant differences in coding precision between degree of syncopation and groove ratings (q < 0.05, FDR-corrected). Error bars indicate SEM.

We quantified this spectro-spatial relationship by fitting the dominant frequency of each vertex to each spatial dimension (x,y,z) and confirmed that this gradient travels conjointly along the antero-posterior (Y) and ventro-dorsal (Z) dimensions, compatible with the localization of the dorsal auditory pathways (Fig. 3b). To investigate if this spectral gradient is specific to music listening or reflects a more generic neurophysiological signature of brain dynamics – i.e., with intrinsic timescales exhibiting a spatial gradient (*36, 37*) along the dorsal auditory pathways–, we performed the same analysis on resting-state data acquired on the same participants (n = 29). We failed to observe a close relation between spectral and spatial dimensions, as indexed by the much less spatially structured pattern of the spectral gradient (Fig. 3c). This dynamic reorganization in the form of a spectral gradient along the dorsal auditory pathways during music listening was confirmed by an individual level estimation of the quality of fit for the two datasets (rm-ANOVA: main effect of dataset: F(1,28) = 54.4, *p* < 0.001; Fig. 3d left).

Further analyses showed that such spectral gradient is a general characteristic of music listening, independent of the specific acoustic, melodic or cognitive attributes of the music. Indeed, we observed that the spectral gradient does not vary across conditions (low, medium, high), neither in shape (Fig. S2a) nor in its quality of fits (Fig. 3d right; repeated-measures ANOVA: main effect of condition: F(2,56) = 0.1, *p* = 0.9). This latter result was robust even at the level of individual melodies (Fig. S2b; repeated-measures ANOVA: main effect of melodies: F(35,980) = 0.8, *p* = 0.7).

Instead, multivariate pattern analyses first revealed that degree of syncopation and groove ratings are both primarily encoded in bilateral and surrounding auditory regions (Fig. 3e-f; q < 0.005 FDR-corrected) in beat-related 2 Hz neural dynamics (Fig. 3h; q < 0.005 FDR-corrected), in agreement with the neurodynamic model. While this neural pattern preferentially encodes the degree of syncopation (Fig. 3g, p < 0.005 uncorrected; Fig. 3h, q < 0.05 FDR-corrected), groove is instead better encoded in the left parietal, supplementary motor, and right motor cortex, at delta (1.3-1.5 Hz) and beta (20-31 Hz and 38-39 Hz) intrinsic neural dynamics (spatial: Fig. 3g, p < 0.005 uncorrected; spectral: Fig. 3h, q < 0.05 FDR-corrected). A complementary regions-of-interest analysis (ROIs estimated from Fig. 3f) confirmed that bilateral auditory regions have a better coding precision for the degree of syncopation at 2 Hz, while more posterior regions better code for groove ratings in delta and (alpha-)beta neural dynamics (Fig. S3; q < 0.05 FDR-corrected; see Methods).

Interestingly, this encoding of groove ratings is adjusted to the spectral gradient of activity along the auditory dorsal pathway (Fig. 3a-b): groove-related delta and beta activity are visible respectively along the inferior portion of the dorsal auditory pathway bilaterally (Fig. 4a; q < 0.005, FDR-corrected) and in more posterior, motor and pre-motor regions (Fig. 4b; q < 0.005, FDR-corrected). Critically, the left sensorimotor cortex acts as the hub coordinating groove-related delta and beta activity through phase-amplitude coupling (Fig. 4c; q < 0.005, FDR-corrected). In this region, delta and beta activity monotonically increase and decrease with groove ratings (Fig. 4d), and the reported coupling is specific to delta (1.3-1.5 Hz) and beta (20-30 Hz) dynamics and is, in particular, not visible for beat-related 2 Hz activity (Fig. 4e). These neural analyses evidence the distributed neural dynamics implicated in the nonlinear transformation from a syncopated stimulus rhythm to the subjective experience of groove, which empirically confirms the results of our neurodynamic model.

**Figure 4.**
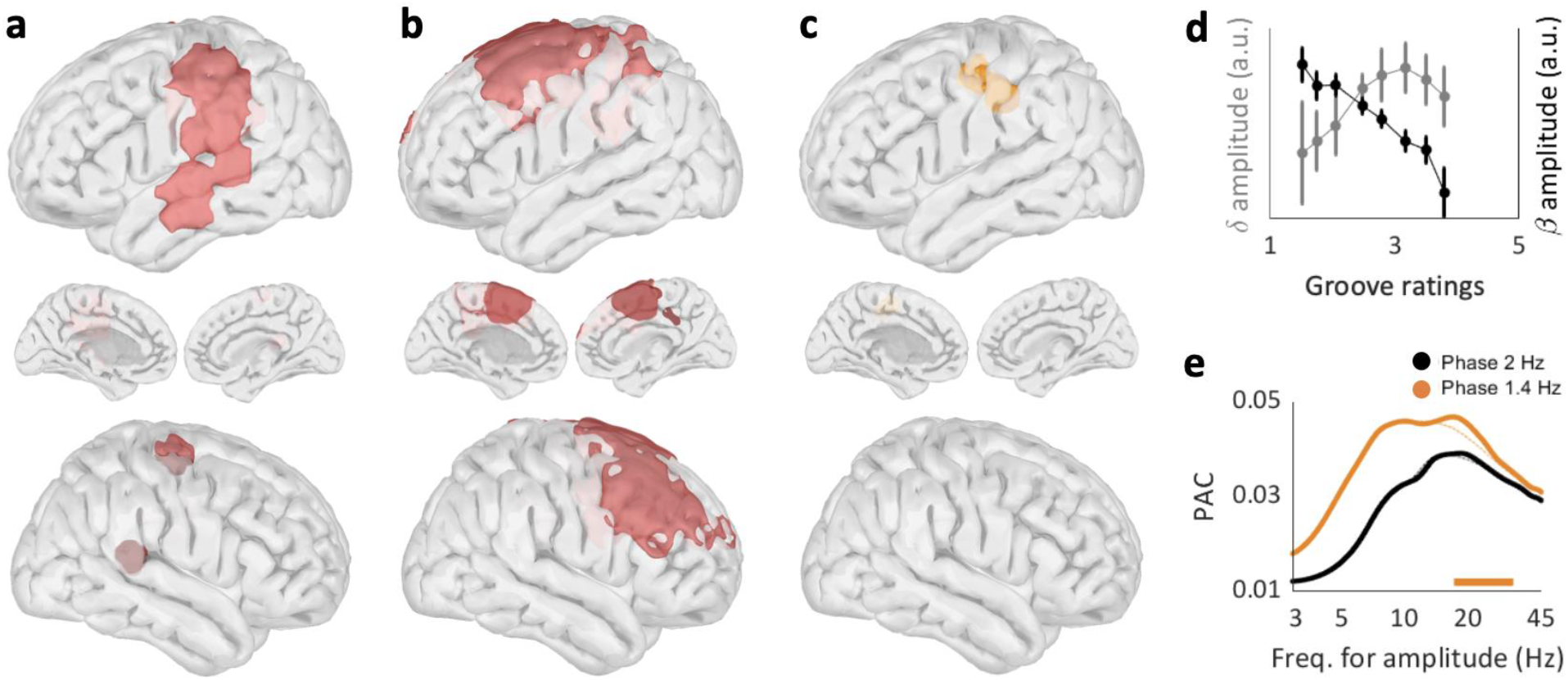
Groove-related neural dynamics in the dorsal auditory pathway. **(a)** Spatial map of neural coding of groove ratings from delta (1.3-1.5 Hz) neural activity. **(b)** Spatial map of neural coding of groove ratings from beta (20-30 Hz) neural activity. **(c)** Spatial map of significant local phase-amplitude coupling (PAC) between delta (1.3-1.5 Hz) phase and any amplitude frequency (3-45 Hz). **(d)** Amplitude of delta (1.3-1.5 Hz; grey) and beta (20-30 Hz; black) neural activity as a function of groove ratings, in the significant PAC cluster (from (c)). Error bars indicate SEM. **(e)** Details of PAC estimates in the significant PAC cluster (from (c); plain lines) or averaged across whole brain (dashed lines), between 1.3-1.5 Hz (orange) or 2 Hz (beat frequency; black) phase and any amplitude frequency (3-45 Hz). The orange horizontal line indicates a significant increase of PAC in the inset cluster compared to whole brain. **(a-e)** All results significant at q < 0.005, FDR-corrected.

## Discussion

Why do humans spontaneously want to move to music? Here, we characterize the computational and neurophysiological bases of this phenomenon by demonstrating that: (i) behavioral motor and neural cortical dynamics principally track the beat of music (Fig. 1e,f); (ii) the pleasurable wanting to move to music -the experience of groove-depends on the temporal regularities present in the music (the degree of syncopation; Fig. 2b-c); (iii) this phenomenon is accounted for by a neurodynamic model and may be described in terms of the timing of auditory events relative to expectancies embodied in oscillations (Fig. 2d-f); (iv) an ascending spectral gradient along the dorsal auditory pathways emerges during music listening (Fig. 3a-d); (v) this gradient supports the encoding of the degree of syncopation in auditory regions and of the experience of groove more dorsally, selectively in delta (1.3-1.5 Hz) and beta (20-30 Hz) neural dynamics (Fig. 3e-h; 4a-b); and (vi) the left sensorimotor cortex acts as a hub coordinating these groove-related neural dynamics (Fig. 4c-e). These results establish the neural dynamics of predictive timing and motor engagement in music listening.

These results extend seminal studies on the quadratic relationship between rhythmic predictability and the experience of groove (*9, 10*). This relationship has been explained using Bayesian inference, with groove arising when precise temporal priors are violated by sensory evidence, here recurring syncopated rhythmic patterns (*5, 38*). Here we provide an alternative dynamical system account (*16*). Mode-locking of neural oscillations to complex rhythms allows the emergence of neural resonance at metrical frequencies. The groove experience is then parsimoniously explained as embodied resonance to a beat and results from the combination of excitatory and inhibitory inputs from two successive layers. This is implemented along the dorsal auditory pathway, which connects the auditory and motor dynamics. The Bayesian and dynamical models converge in the sense that groove may be described as affordance for movements that interact with the stimulus temporal structure, relying on the timing of auditory events relative to expectancies. The neurodynamic model further specifies temporal expectations as being the consequence of neural resonance, i.e. being embodied in oscillations, and hence provides a physiologically plausible model to predictive timing (*7, 18, 39, 40*) and the experience of groove (*5, 15*).

That the motor system contributes to auditory perception is also supported by human psychophysics and neuroimaging data, in particular when listening to rhythmic auditory streams (*19, 25, 26, 29*). Here we extend these findings to an ecological music listening situation and also show that the dissociation between auditory and higher-level cognitive functions in the dorsal auditory pathway is subtended by a spectral dissociation. This anatomo-spectral gradient could be instrumental in structuring the information flow. Our finding extends a few recent reports of the presence of a spectral gradient along ventral auditory or visual dorsal cortical pathways (*36, 37*) and provide clear evidence of a tight relation between the anatomical and dynamical dimensions of the brain. Whether this gradient acts as a support for information transfer, or directly encodes information remains to be investigated, but our results indicate that it does not code for the sensory (degree of syncopation) or cognitive (groove ratings) investigated variables.

During music listening, this gradient is organized around the left sensorimotor cortex, a hub implicated in feedback signaling during auditory perception (*19, 41*). This region might correspond to area 55b, a proposed keystone of sensorimotor integration critical in both music (*42*) and speech (*43, 44*) perception. This hub coordinates ventral delta (1.4 Hz) and dorsal beta (20-30 Hz) intrinsic neural dynamics and is hence constitutive of the emergence of audio-motor coupling. While beta dynamics (∼12–30 Hz) are typically associated with movement planning and execution, the specific role of delta (1.4 Hz) activity in the auditory dorsal pathway corresponds to the optimal rate for auditory temporal predictions and defines the auditory temporal attention capacity (*27*). This support a model that considers neuronal oscillations as intrinsic dynamical mechanisms capable of embodying neural computation (*45, 46*). In conclusion, we show how interacting neural dynamics along the dorsal auditory pathway effect the spontaneous emergence of the pleasurable wanting to move during music listening.

## Acknowledgments

We warmly thank Aurélie Ponz and Sophie Chen for their contribution to MEG data acquisition and Christian Bénar for discussions on the paradigm. Research supported by grants ANR-20-CE28-0007 (BM), ERC-CoG-101043344 (BM), ANR-21-CE28-0010 (DS), ANR-16-CONV-0002 (ILCB) and the Excellence Initiative of Aix-Marseille University (A*MIDEX).

## Funding

ANR-20-CE28-0007 (BM)

ERC-CoG-101043344 (BM)

NR-21-CE28-0010 (DS)

ANR-16-CONV-0002 (ILCB) and the Excellence Initiative of Aix-Marseille University (A*MIDEX).

SPARK Award from University of Connecticut (EWL)

## Author contributions

Conceptualization: BM, DS

Methodology: AZ, EWL, DS, BM

Investigation: AZ, BM

Visualization: AZ, BM

Funding acquisition: BM

Project administration: BM

Supervision: BM

Writing – original draft: AZ, BM

Writing – review & editing: AZ, EWL, DS, BM

## Data and materials availability

Numerical data supporting this study will be available on GitHub upon publication: https://github.com/DCP-INS/Groove/.

## Materials and Methods

### Participants and stimuli

#### Participants

66, 30 and 15 participants (age range: 19-71 years; 77 % females) were recruited for the online, Magnetoencephalography (MEG) and control tapping experiments. All experiments followed the local ethics guidelines from Aix-Marseille University. Informed consent was obtained from all participants before the experiments. All had normal audition and vision and reported no history of neurological or psychiatric disorders. We did not select participants based on musical training and a short survey made at the end of the experiment informed us that none of them were professional musicians. Participants were financially compensated for their time during the MEG experiment.

#### Acoustic stimuli

A professional musician composed in MIDI format 36 melodies lasting 8 seconds each (stimuli are available on github.com/DCP-INS/Groove). All had the same strictly periodic drum beat at 2 Hz combined with a specific bass melody. To vary the level of rhythmic predictability while minimizing other acoustic variations, three variants were derived from each of 12 original melodies, using an ascending degree of syncopation ((the amount of syncopation is inversely proportional to the rhythmic predictability). The bass melodies were maximally matched between the three variants of the same original melody, with the number of notes per measure, their pitch and their order being identical or closely matched. Other musical characteristics (volume, timbre, etc.) were kept constant. This procedure resulted in the creation of three conditions, reflecting low (black), medium (grey) or high (light grey) degree of syncopation (Fig. 1). Syncopation was defined as the appearance of a beat on a metrically weak accent preceding a rest on a metrically strong accent and quantified after Longuet-Higgins and Lee (*47*). Finally, the songs were recorded in stereo with a sampling rate of 48 kHz and a bit-depth of 24-bit.

### Experimental designs and data acquisition

#### Experimental design of the online experiment

Participants were invited to visit a web page to take part in the survey, hosted by PsyToolkit. After completing the questionnaire, participants were invited to start the experiment. They were first prompted to use headphones or earphones and given the opportunity to pre-evaluate the output volume in order to adjust them comfortably. Then, the experiment consisted of a listening task in which each of the 36 melodies was presented binaurally once to participants, in a randomized manner. After stimulus offset, participants reported on a keyboard the associated level of groove, defined as the extent to which they wanted to move to this music (*9–15*). They had 60 seconds to answer. A Likert scale between 1 and 7 was used. Instructions were visually displayed on a mid-grey background on a screen computer. During each trial participants had to fixate a cross, located at the center of the screen. The online experiment lasted approximately 10-15 minutes.

#### Experimental design of the control tapping experiment

The experiment was performed in the laboratory, using the Psychophysics-3 Toolbox and additional custom scripts written for MATLAB (The MathWorks). Trials consisted of a tapping task in which participants were asked to reproduce, with their indexes on the computer keyboard, the rhythm of the dance step they would naturally produce when listening to the melodies. Each of the 36 melodies was presented binaurally once to participants, in a randomized manner, at a comfortable hearing level via headphones. Instructions were visually displayed on a mid-grey background on the screen laptop situated at a viewing distance of around 50 cm. On each trial, participants had to fixate a cross, located at the center of the screen.

#### Experimental design of the MEG experiment

The experiment consisted of a listening task in which melodies were presented binaurally to participants, in a randomized manner. Participants were requested to stay completely still while they were listening to the melodies. Each original melody was duplicated, while maintaining the beat structure, which resulted in 16-second long melodies. This allowed us to optimize the signal-to-noise ratio of the MEG response. Moreover, the experiment was composed of 4 blocs. In each bloc, each of the 36 duplicated melodies was presented binaurally once to participants, in a randomized manner (144 trials in total). After stimulus offset, participants had 4 seconds to report on a keyboard the associated level of groove on a scale between 1 and 5. The experiment lasted approximately 48 minutes.

#### Experimental design of the resting state experiment

Participants performed two 4-minute eyes-open resting-state sessions, at the beginning and at the end of the MEG experiment.

#### MEG data acquisition

MEG data were acquired at the Epileptology and Cerebral Rhythmology Unit from the La Timone hospital, APHM, Marseille (France), using a 4D Neuroimaging™ 3600 whole head system (4-D Neuroimaging, San Diego, CA, USA) composed of 248 magnetometers, at a sampling frequency of 2034.51 Hz EOG and ECG channels, one audio and five response buttons (LUMItouch optical response keypad) were recorded simultaneously and synchronized with the MEG signal. Presentation software was used for stimulus delivery and experimental control during MEG acquisition. Auditory stimuli were presented binaurally at a comfortable hearing level through insert earphones (E-A-RTONE 3A, Aero Company). Instructions were visually displayed on a mid-grey background on a screen computer situated at a viewing distance of around 50 cm. On each trial participants had to fixate a cross, located at the center of the screen, to get a visual constant stimulation. Location of the participant’s head with respect to the MEG sensors was recorded both at the beginning and end of each session to potentially exclude sessions and/or participants with large head movements. However, none of the participants moved >3 mm during all sessions.

#### MRI data acquisition

For volume MEG source analysis (i.e., the projection of the MEG sensor data onto the full brain volume), a T1-weighted MRI acquisition of the brain was obtained from each participant (1-mm isotropic voxel resolution).

### Data analyses

#### Timing of motor acts in the control tapping experiment

To investigate the dynamics of occurrence of the motor events produced by participants during the listening of the melodies, we estimated the instantaneous frequency of their finger taps by computing the inverse of the inter-tap interval (in Hz; i.e. 1/(tn-tn-1)).

#### Spectral decomposition of the acoustic stimuli

To estimate the temporal envelope of each melody, the sound signal was decomposed into 32 narrow frequency bands using a cochlear model, and the absolute value of the Hilbert transform was computed for each of these narrowband signals. The broadband temporal envelope resulted from the summation of these absolute values and was used as the acoustic signal for all subsequent analyses. Using a fast Fourier transform, we then decomposed the acoustic signal of each melody from 1 to 9 Hz, to obtain the acoustic temporal modulation spectrum (i.e., the spectrum of the temporal envelope).

#### MEG data preprocessing

Preprocessing was performed with Brainstorm (*48*), following good-practice guidelines (*49*). Briefly, we removed electrical artefacts using notch filters (at 50 Hz and its first three harmonics), slow drifts using high pass filtering (at 0.3 Hz), and eye blink and heartbeat artefacts using source signal projections. Data were split into 20-second trials, from -2 to +18 seconds relative to stimulus onset. MRI volume data were segmented with Freesurfer and transformed in MNI space. A template source grid covering the entire brain volume was created on the default anatomy (10 mm resolution) and projected to individual anatomies to be used for the individual source-reconstruction procedure. We computed individual MEG forward head models using the overlapping-sphere method (volume), and source imaging using dSPM (v. 2016, median eigenvalue) onto preprocessed data, all by using default Brainstorm parameters. We obtained 1673 source volumes (i.e., vertices), each composed of three orientations (x,y,z). The procedure also included an empirical estimate of the variance of the noise at each MEG sensor, obtained from a 2-min empty-room recording done at the beginning of each scanning session. One participant was excluded from subsequent analyses (hence n = 29) as we failed to detect proper auditory responses.

#### Spectral decomposition of the MEG data

For both channel-level and source-level MEG data, trial-by-trial time-frequency decomposition was conducted in a range of 100 frequencies, logarithmically spaced from 1 to 100 Hz. Morlet wavelet transform was applied to the data using the Brainstorm (Matlab) function bst_timefreq with parameter Method = ‘morlet’, central frequency Morlet_Fc = 1 and time resolution Morlet_FwhmTc = 3.

#### Normalized power spectrum

For each vertex, trial and participant, we estimated the power spectrum by time-averaging the spectrally decomposed signals in the [0.5-16] seconds range. 1/f aperiodic component was removed by z-scoring the data across voxels for each frequency. We hence obtained, for each frequency, the distribution of power across voxels, centered around zero.

#### Modelling of spectral gradient

We fitted the source-reconstructed spatial distribution of the MEG power spectrum, both at the group- and at the individual-levels, for both the MEG experiment and resting-state datasets. We approximated the data with a five-order polynomial function, fitting the dominant frequency of each vertex to each spatial dimension (x,y,z) and projecting the results in the 3D space.

#### Multivariate pattern analysis on channel-level MEG data

Multivariate pattern analyses were conducted by capitalizing on the spatial patterns of the MEG power signal, i.e., spectrally decomposed and time-averaged (0.5-16 seconds, to exclude the stimulus-onset period), for each participant, each neural frequency and each regressor (degree of syncopation, groove ratings, acoustic temporal modulation spectrum). We used a cross-validated multivariate linear encoding model to estimate the spatial MEG patterns *ŵ* of a specific data associated with each stimulus characteristic X. For each cross-validation fold (n = 10, interleaved, see (*50*)), we defined the spatial MEG patterns *ŵ* on the training set by regressing—in a ridge sense (ridge α parameter set at 2) —each z-scored MEG feature *Z*_*t*rain_ (248 channels in total) against the stimulus characteristic *X*_*train*_ across stimulus exemplars (n = 130 for each cross-validation fold), by solving 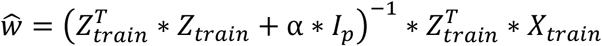, where *I*_*p*_ is the *p*p* identity matrix with *p* corresponding to the number of MEG channels (248 in total). We then projected the MEG data on the test set *Z*_*test*_, on the dimension defined by the coding weights *ŵ* to obtain neural predictions of the stimulus characteristic *X*_*test*_ for each epoch of the test set (n = 14 for each cross-validation fold). After applying this procedure for each cross-validation fold, we computed the linear Pearson’s correlation coefficient between neural predictions 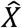 and ground-truth values X of the stimulus characteristic. The coding precision metric reported in the main text corresponds to the Fisher transform of the correlation coefficient, which is approximately normally distributed, such that we could compute standard parametric statistics at the group level.

#### Searchlight analysis on source-level MEG data

We conducted searchlight-based multivariate pattern analyses on the entire power signal of the reconstructed volume sources (vertices), for each participant and each regressor (degree of syncopation and groove ratings). The searchlight procedure was applied to each vertex position by using as features the entire power spectrum of the current vertex and its fifty closest neighbors (in terms of Euclidean distance).

#### Multivariate pattern analysis on regions-of-interest

We defined five regions-of-interest of 20 vertices each based on the contrast between the decoding of groove ratings and degree of syncopation at the source level (Fig. 3g and S2). The 20 vertices surrounding (in terms of Euclidean distance) the maximally significant vertex where selected. We then conducted multivariate pattern analyses on the power signal, for each participant, each frequency, each regressor, and each region-of-interest.

#### Phase-Amplitude Coupling

We estimated phase-amplitude coupling over time (0.5-16 seconds) and melodies for each source-reconstructed vertex, between the phase at 1.4 or 2 Hz, and the amplitude between 3 and 45 Hz (*51*).

#### Statistical Procedures

All analyses were performed at the single-subject level and followed by standard parametric tests at the group level (e.g., two-tail paired t tests, two-tail t tests against zero, repeated-measure ANOVAs). The type 1 error rate arising from multiple comparisons was controlled for using False Discovery Rate (FDR) correction over the dimensions of interest (i.e., time, vertices, frequencies), using the Benjamini–Hochberg step-up procedure.

#### Neural network model

We implemented a canonical model for Gradient Frequency Neural Networks based on the GrFNN Toolbox (for more information please see on musicdynamicslab.uconn.edu/home/multimedia/grfnn-toolbox/ and (*16, 17, 32, 33, 45, 46*)). The aim of this modelling approach was twofold. We first investigated whether neural resonance can explain the experience of groove during music listening, and specifically whether a neurodynamic model can capture the nonlinear relationship between degree of syncopation and groove ratings (Fig. 2b-c). Our secondary goal was to understand which neurodynamic mechanisms may underlie the experience of groove.

Our model is composed of three 1-dimensional networks of nonlinear oscillators, tuned to different natural frequencies. Such networks are conceptually like banks of band-pass filters, except that they consist of nonlinear oscillators rather than linear resonators. Network elements are canonical Hopf oscillators, a fully expanded canonical model for excitation-inhibition oscillations near a Andronov-Hopf bifurcation (*17, 45, 52*). As a generic model of excitation-inhibition oscillations, the canonical model represents the firing rates of interacting excitatory and inhibitory neural subpopulations as sinusoidal oscillations in the complex plane. The oscillators of each network are tuned to a range of distinct frequencies spanning the delta - theta frequency range, and stimulated with time-varying acoustic signals given by:

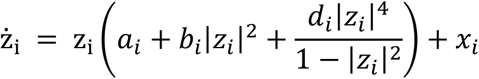

where *z*_*i*_ is the complex-valued state of the ith oscillator in the network (subscript i = 1, …, N), a_i_ = α_i_ + iω_i_, b_i_ = β_1i_ + iδ_1i_, d_i_ = β_2i_ + iδ_2i_(α_i_, ω_i_, β_1i_, δ_1i_, β_2i_, δ_2i_ ∈ R; i denotes the imagery unit), and *x*_*i*_ is the sum of input terms. The parameters α_*i*_, β_1i_ and β_2i_ determine the intrinsic dynamics of the i-th oscillator, where α_*i*_ is the bifurcation parameter; ω_*i*_ is its natural frequency; δ_1i_ and δ_2i_ determine the dependence of intrinsic frequency on amplitude. The input to the i-th oscillator *x*_*i*_ can include both an external signal *s*_*i*_(*t*) corresponding to the onsets of rhythmic auditory stimuli and coupling from other oscillators,

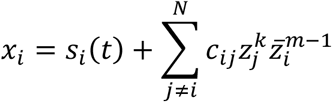

where *c*_*ij*_ is the coupling coefficient (*17*), and k:m is the mode-locking ratio (i.e., 1:1, 2:1, 3:1, etc.).

For the first (auditory) layer we set the parameters as α = 0.0001, β_1_ = 0, β_2_ = −3. Thus, each oscillator in this network produces a low amplitude intrinsic oscillation at its own intrinsic frequency that can be entrained by an external rhythmic stimulus (*32, 53, 54*). The second (motor planning) layer parameters were chosen as α = −0.8, β_1_ = 4, β_2_ = −3. In this parameter regime, each oscillator was bistable (or double limit cycle) regime, exhibiting a both stable limit cycle at zero, and a stable limit cycle at a higher amplitude. Oscillators in this network begin at rest, and when it receives a signal that is strong enough and long enough, it can jump to a stable limit cycle. The third (groove) layer has the same parameters as layer two. The remaining parameters were all set to zero: δ_1_ = 0, δ_2_ = 0. All the layers were based on 321 oscillators of internal frequency ω_*i*_ which span a range between 0.375 and 12 Hz (inclusive of the delta - theta frequency range) according to a log scale.

In this network, connections are learned according to the Hebbian rule (*46, 55*):

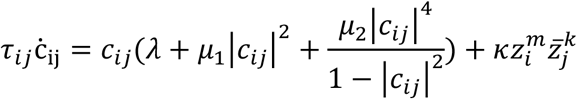

To make the dynamical interactions as transparent as possible, we designed a feedforward model with a minimal connection topology. Layer 1 is connected only to the auditory input, represented by the onset of the notes in the stimuli. We assumed learned multifrequency connections (black arrows in Fig. 2d) between layers 1 and 2. Initial connection strengths were chosen such that each frequency is connected to its 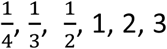 and 4^th^ harmonics. We then fixed the learning parameters as λ = −1, μ_1_ = 4, μ_2_ = −2.2, κ = 0.2 (*46*). We then assumed fixed 1:1 excitatory connections from layer 2 to 3 with a coupling strength fixed as *w* = 0.8, and fixed 1:1 inhibitory connections from layer 1 to 3 (red arrow in Fig. 2d) with a coupling strength fixed as *w* = -0.7. Thus, oscillators in layer 3 received in-phase stimulation from the motor planning network, and anti-phase stimulation from the auditory network. This means that the input to layer 3 was the difference between the time-dependent oscillations of layer 2 (pulse/meter) and layer 1 (auditory rhythm).

We ran the model as follow: The MIDI^1^ representation of each melody was used to provide a clear estimate of the note onsets. Note onsets were encoded as a sequence of continuous time onset pulses and transformed into a complex-valued signal using a Hilbert transform^2^. The model entrained to the onsets for the entire duration of the melodies (16 sec). As with the data from the human experiment, we removed the initial (evoked) response of the model (here, the first 2 sec). We then computed the mean field of each network individually (the mean of all oscillators in the network; see (*16*)). Finally, we computed fft of the mean field (for seconds 2-16 of the stimulus) and extracted the amplitude of the response at 2 Hz for each layer of the model, for each melody. We repeated this procedure 29 times (similar to the number of participants in our MEG experiment) to obtain a robust estimate of the model’s response^3^. Overall, this procedure produced an estimate of the amplitude of the time-averaged 2 Hz oscillations in each of the three network layers, in response to each of the 36 melodies.

## Supplementary Text

### Supplementary Results

#### Modelling the relation between groove ratings and degree of syncopation

We fitted at the individual level the relation between groove ratings and degree of syncopation. We fitted both a linear and quadratic model, and compared their goodness-of-fit, by using the adjusted r-squared values. For the online experiment (n = 66), we obtained an average adjusted r-squared of 0.14 and 0.37 across melodies, for the linear and quadratic functions. The quadratic model significantly outperformed the linear one (t(65) = 13.1; *p* < 0.001). For the MEG experiment (n = 29), we obtained an average adjusted r-squared of 0.21 and 0.49 across melodies, for the linear and quadratic functions. The quadratic model significantly outperformed the linear one (t(28) = 11.8; *p* < 0.001). At the group level, the inverse U-shape profile is very well approximated with a quadratic function for both the online (adjusted r^2^(33) = 0.73) and MEG (adjusted r^2^(33) = 0.67) experiments.

#### Correlation of the degree of syncopation and groove ratings with the neural network model outputs

We first correlated the degree of syncopation with the time-averaged 2 Hz amplitude of the dynamics of each layer of the neurodynamic model. Layer 1 strongly linearly correlates with the degree of syncopation (r^2^(34) = 0.85; *p* < 0.001), which is not the case for layer 2 (r^2^(34) = 0.12; *p* = 0.04) or layer 3 (r^2^(34) = 0.14; *p* = 0.02). Second, we correlated groove ratings, averaged across participants, obtained either from the online (n = 66) or the MEG (n = 29) experiment. Groove ratings mostly linearly correlated with activity from layer 3 (online: r^2^(34) = 0.66; *p* < 0.001; MEG: r^2^(34) = 0.57; *p* < 0.001), but to a far less extent with activity from layer 1 (online: r^2^(34) = 0.34; *p* < 0.001; MEG: r^2^(34) = 0.38; *p* < 0.001) or layer 2 (online: r^2^(34) = 0.19; *p* = 0.008; MEG: r^2^(34) = 0.13; *p* = 0.03). Our results show that a high groove experience reflects strong resonance in layer 2 (pulse/meter) at frequencies that are weak and/or anti-phase in the stimulus (and in layer 1), while difficulty or inability to perceive a pulse (the absence of *expectancy* in layer 2) is characteristic of high syncopated melodies (Fig. S1b).

**Figure S1.**
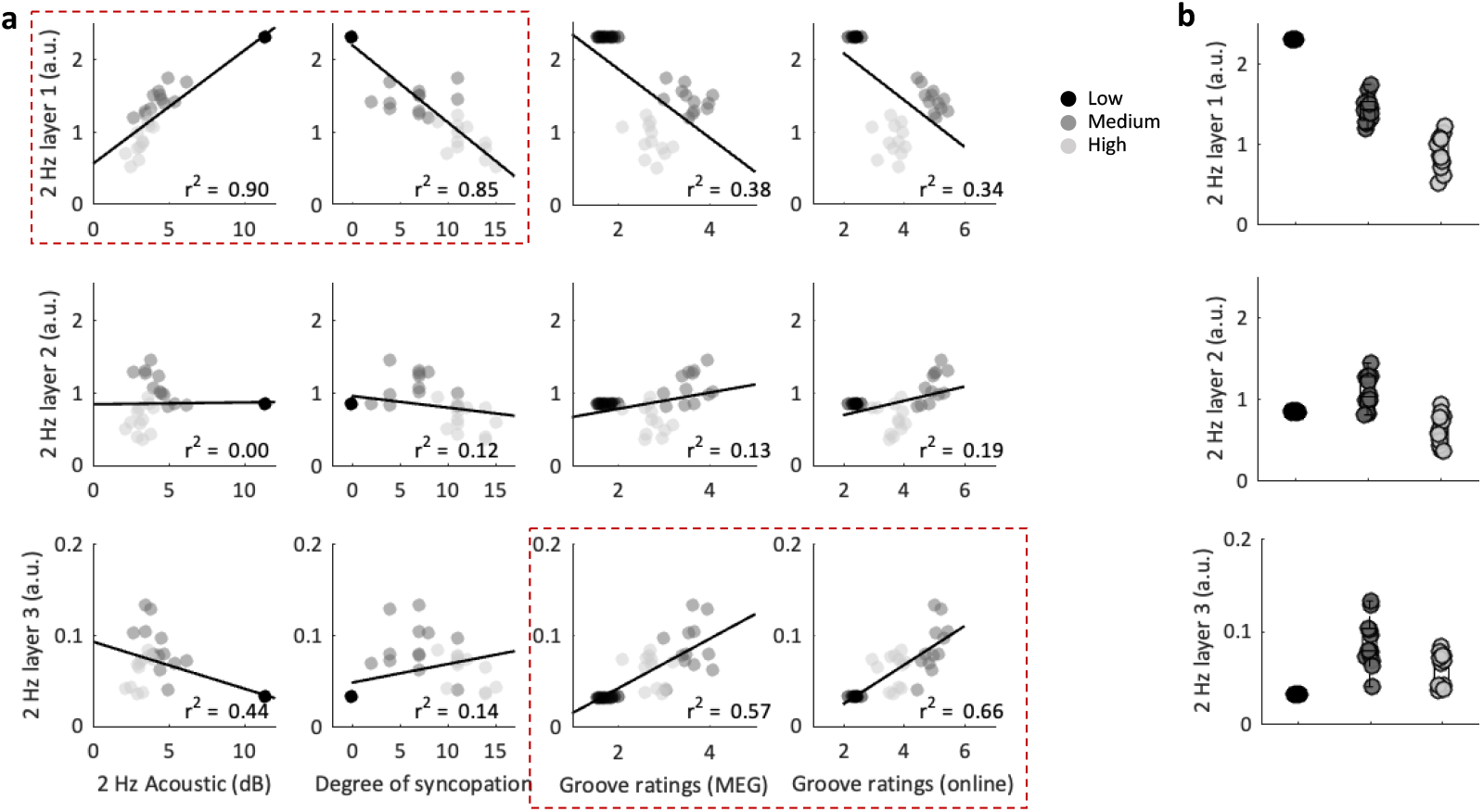
Neurodynamic model. **(a)** Inter-melody correlation between the time-averaged 2 Hz amplitude of each layer of the neurodynamical model (1 to 3; lines, y-axes), as a function of 4 paradigmatic variables (2 Hz acoustic, degree of syncopation, MEG groove ratings, or online groove ratings; columns, x-axes). Data were approximated with a linear function. Pearson’s r-squared is reported. Strongest correlations are highlighted with red rectangles. **(b)** Amplitude of the time-averaged 2 Hz oscillations in the three network layers, in response to each of the 36 melodies, for each condition (low, medium, high). Shades of grey indicate the conditions. Individual points indicate melodies (n = 36).

**Figure S2.**
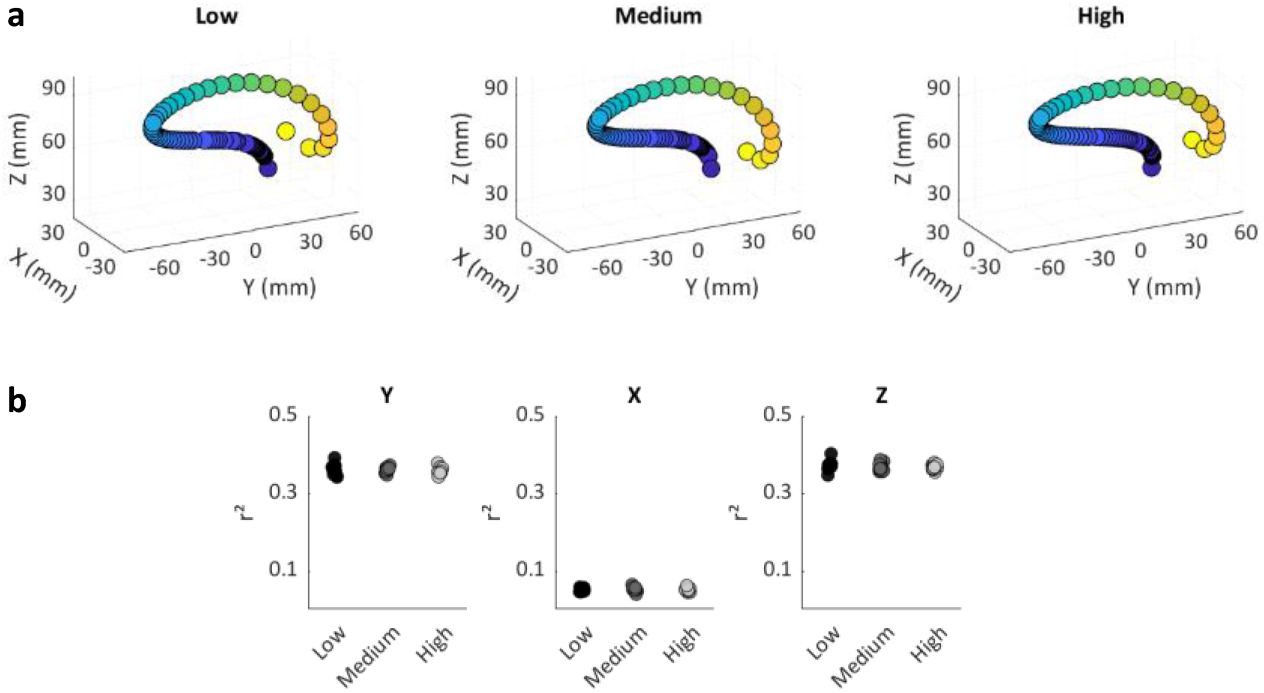
Dominant frequency across the brain volume in the three melodic conditions (low, medium, high). Frequency range 1-45 Hz (after removal of the 1/f decay of the neural power spectrum). **(a)** For each condition, data were approximated at the group-level with a polynomial function, independently for each dimension (X, Y, Z) of the MNI space. **(b)** Comparison of the quality of fits (r^2^), estimated at the individual level, of the 36 melodies grouped per condition (low, medium, high) and plotted for each dimension (Y, X, Z) of the MNI space.

**Figure S3.**
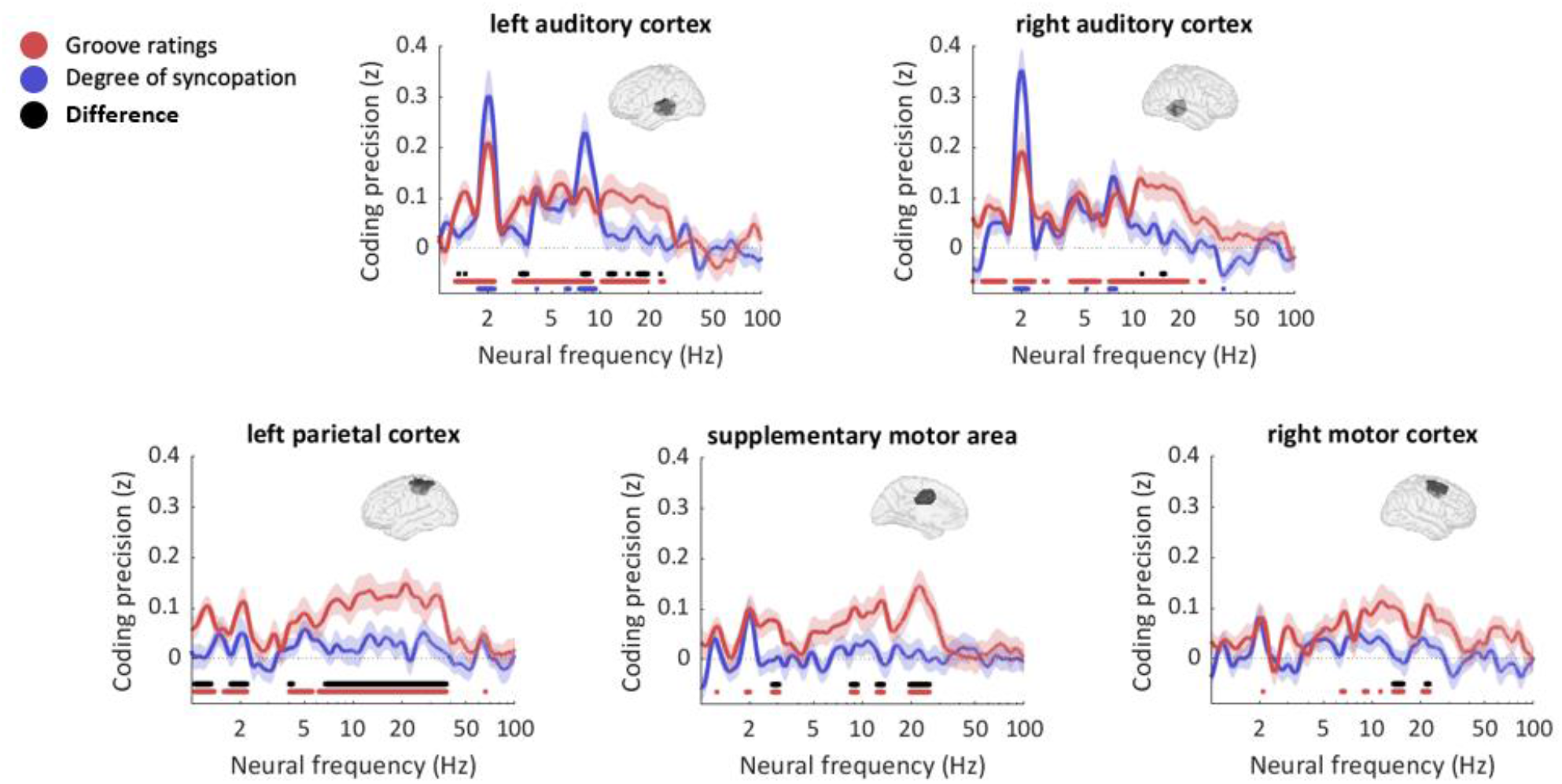
MEG experiment: Spectral coding of degree of syncopation and groove ratings in the regions-of-interest. Spectrum of neural coding of the degree of syncopation (blue) and groove ratings (blue), for each of the regions-of-interest (Fig. 3g). Red and blue horizontal lines indicate frequencies with significant coding values (q < 0.005, FDR-corrected). The black line indicates frequencies with significant differences in coding precision between degree of syncopation and groove ratings (q < 0.05, FDR-corrected). Error bars indicate SEM. Inset brains indicate the spatial localization of each region-of-interest.

## Footnotes

1 Musical Instrument Digital Interface

2 We could equally well have used a real-valued signal. However, the complex input is more natural for this model.

3 The network dynamics were deterministic, but initial conditions were chosen at random, so the network output for each run of the same stimulus differed somewhat.

